# A Mechanistic Model of Caspase-mediated Apoptosis Signaling Reveals Regulators of Drug Resistance in Prostate Cancer Cells

**DOI:** 10.64898/2026.03.09.710645

**Authors:** Diamond S. Mangrum, Stacey D. Finley

## Abstract

Anticancer drug resistance remains a major barrier to effective treatment because both intrinsic and acquired resistance mechanisms can suppress apoptotic cell death. The growing incidence of castration-resistant prostate cancer highlights the need for therapeutic strategies that remain effective, independent of hormonal signaling. Here, we developed a parsimonious, data-calibrated ordinary differential equation model of caspase-mediated apoptosis signaling in prostate cancer. Rather than representing the full complexity of the apoptotic network, the model incorporates a reduced set of key apoptotic regulators required to capture experimentally-observed apoptotic responses. Model parameters were estimated using published data from three pro-apoptotic agents (Tocopheryloxybutyrate, Narciclasine, and Celecoxib), and global sensitivity analysis was used to identify determinants of treatment response and resistance. The model accurately reproduced experimental observations and revealed that variability in specific signaling components can produce substantial differences in response to pro-apoptotic drug treatment. Simulations identified key regulators of apoptotic commitment, predicted conditions under which resistance may be overcome through combination therapies, and suggested candidate mechanisms underlying heterogeneous responses among cancer cells. Together, these findings demonstrate that a reduced mechanistic framework can generate biologically meaningful insight into apoptotic drug response while identifying testable hypotheses regarding therapeutic vulnerabilities. This work establishes a quantitative platform for investigating resistance mechanisms and guiding future experimental studies.

## Introduction

Despite advances in oncology, treatment failure persists across all major anticancer strategies, including chemotherapy, targeted therapy, radiation, and immunotherapy. One 2020 study indicates that nearly half of United States cancer patients eligible for cytotoxic chemotherapy do not respond to treatment.^1^ Targeted therapies and immunotherapy are also susceptible to physiological resistance, as highly adaptive cells acquire mutations that allow them to evade therapeutics over time.^2^ Due to its emergence through both intrinsic and acquired mechanisms, regulated by distinct cellular pathways, drug resistance is difficult to overcome for several reasons.

First, the immune system is a complex network designed to safeguard the body against harmful, foreign, and threatening invaders. While both the innate and adaptive immune systems are highly successful in invading harmful perturbations, this same capability can be used inversely against anticancer drugs.^3^ Second, therapeutic agents may target components of the apoptotic pathway that are not the primary source of dysfunction, allowing pro-survival signaling to persist. Third, heterogeneity across tumors, within individual tumors, and within patients’ immune systems introduces an additional layer of complexity that contributes to natural resistance. Alongside this issue, in some patients, cancer cells fail to respond to apoptotic induction because intracellular drug concentrations do not reach levels required to inactivate the target.^4^ Given these causes, a large portion of research focuses on strategies to overcome natural, or intrinsic, drug resistance by developing strategies that either combine drugs, curate the treatment to fit patient profiles, or design mechanisms that strengthen the antitumor immune state of the tumor microenvironment.^5^

All of these treatment strategies can be strengthened by the effective removal of damaged, infected, or cancerous cells, which is initiated by apoptosis. Apoptosis-targeted therapies aim to bolster the critical innate biological process that keeps the homeostatic balance of cells in healthy tissues. Unlike some anticancer approaches that can result in associative damage to surrounding tissues and the release of inflammatory substances (i.e. radiation, chemotherapy, surgical removal, etc.), apoptotic signaling prioritizes a non-reactive response.^6,7^ Without apoptosis, healthy cells would proliferate uncontrollably to form large masses that disrupt normal physiological function. This role in preventing overproliferation is why apoptotic evasion is considered one of the nine hallmarks of cancer.^8^

Apoptosis can be regulated through both its intrinsic and extrinsic signaling pathways. However, crosstalk between these two pathways can drive both tumor progression and death in hormone-dependent cancer types.^9^ For example, apoptotic regulators such as caspases mediated by death inducing signaling complexes (DISC), act variably in response to a range of ligand-receptor binding interactions.^10^ Thus, it is unclear whether apoptotic mediators are directly responsible for the cancer cell-killing effects of many pro-apoptotic drugs and whether sustained signaling of apoptosis could lead to unintended effects.^11,12^ Motivated by this gap, we leverage mechanistic modeling, which enables *in silico* perturbation of individual components, to determine their causal roles in apoptosis execution in the PC3 castration-resistant prostate cancer cell line. Prostate cancer, affecting 1 in 8 men, could benefit from more informed approaches that enhance apoptotic potential, particularly given that many current treatments fail to adapt to the transition from hormone-sensitive to hormone-insensitive prostate cancer cells.^13^ As an example, androgen deprivation therapy (ADT), used to shrink prostate tumors, often causes prostate cancer cells to evolve into castration-resistant prostate cancer (CRPC).^14^ ADTs rely on antiandrogens, such as bicalutamide and enzalutamide, to block the formation of androgen receptor complexes. However, once CRPC is reached, prostate cancer can continue to progress despite low androgen levels. As an exemplary case, one study revealed that a third of patients that underwent enzalutamide treatment and up to one third of patients that underwent abiraterone (another antiandrogen) treatment exhibited intrinsic resistance to their respective drugs.^15^

Optimizing the role of the combined (intrinsic and extrinsic) apoptosis pathways for prostate cancer treatment represents a promising next step, since apoptosis signaling operates independently of androgens. To aid in optimizing apoptosis-targeting prostate cancer treatment, in this work we develop and analyze an ordinary differential equation (ODE) model of caspase-mediated apoptosis that integrates key proteins from both the extrinsic and intrinsic pathways. The model simulates the dynamics of three separate prostate cancer drug-response datasets and quantifies each drug’s effect on apoptotic potential. Building on our prior Fas-mediated apoptosis model^10^ and a published modular caspase activation model^16^, we extend these frameworks to develop a quantitative model that provides quantitative insight into why some prostate cancer cells fail to respond to pro-apoptotic drugs. By doing this, we address two central questions: (1) Can caspase-mediated apoptosis dynamics be accurately captured in response to three separate prostate cancer pro-apoptotic drugs? (2) Which features of the integrated apoptosis pathway represent viable targets for overcoming treatment resistance?

## Methods

To answer these two central questions, we constructed an *in silico* model in MATLAB that simulates caspase-mediated apoptosis signaling in individual prostate cancer cells using ODEs deduced using the law of mass action. The ODE model captures the continuous, time-dependent dynamics and nonlinear kinetics of molecular species, simulating each *in silico* cell for 48 hours to maintain relevance to *in vitro* experiments and to encompass the timescales reflected in the calibration data.^17^ We began with the extrinsic arm modeled in prior work, composed of eleven ODEs, representing distinct species on the extrinsic pathway and the following dynamics: receptor degradation, binding dissociation, and complex internalization, promoting caspase activation.^10^ Then, we added 10 additional molecular species that are characteristic proteins on the intrinsic apoptosis arm. Lastly, apoptosis signaling was partitioned into three core regulatory modules: the death-inducing signaling complex (DISC), mitochondrial-activated caspase (MAC), and apoptosome, building on the modular caspase activation model developed by Harrington and co-workers.^16^ This modular structure enables systematic representation of crosstalk between the extrinsic and intrinsic pathways while preserving biological interpretability. In total, the resulting model is composed of 21 molecular species, 21 ordinary differential equations (ODEs), and 14 reaction rate expressions. To address the first central question of this work, *whether apoptosis dynamics can be accurately captured in response to three prostate cancer pro-apoptotic drugs*, we adapted our previously established non-tumor-specific model to a prostate cancer-specific drug-response model. This adaptation was performed using an integrated pipeline of ImageJ data extraction, particle swarm optimization (PSO), and extended Fourier Amplitude Sensitivity Testing (eFAST) (**Figure 1**).

**Figure 1:**
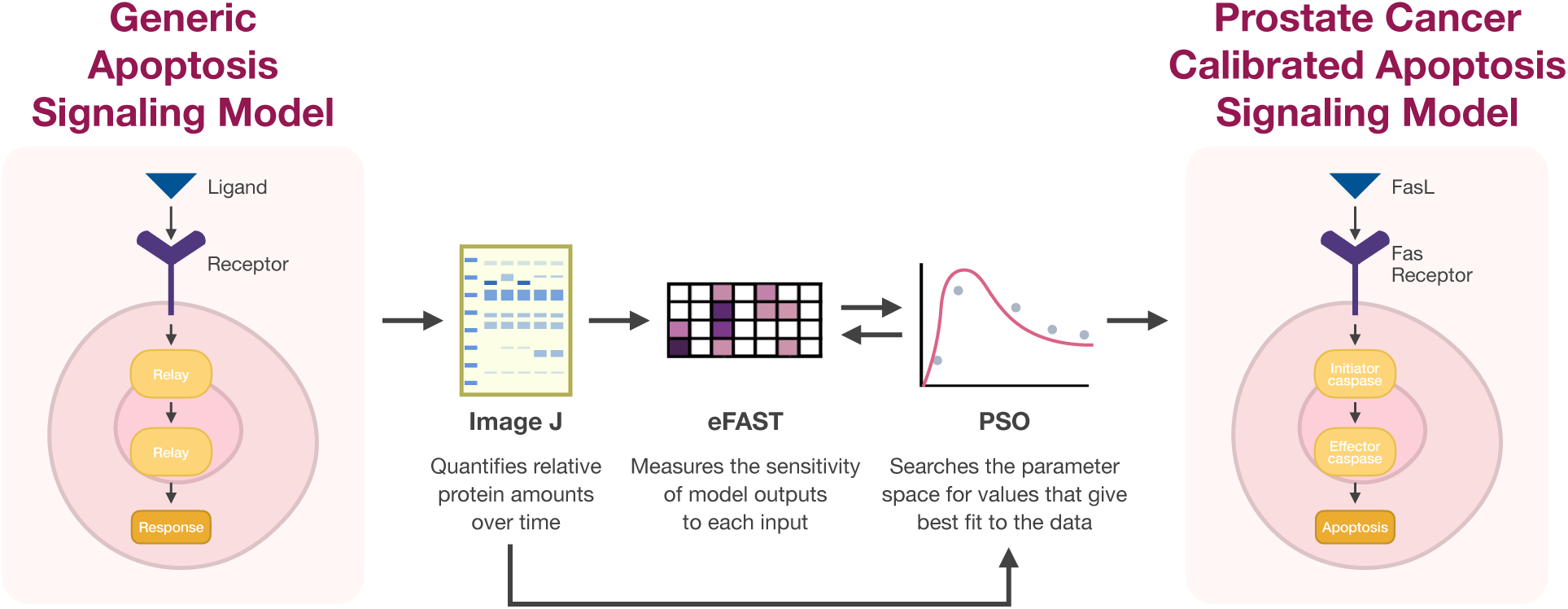
Computational workflow for model calibration. The computational method workflow for calibrating the model to three separate pro-apoptotic drugs acting upon the PC3 cell line: data extraction using ImageJ, sensitivity analysis using the eFAST method, and parameter estimation using particle swarm optimization (PSO).

### Overview of Generic Model of Apoptosis Signaling

The signaling pathway is stimulated extrinsically by FasL binding to its receptor, creating DISC. DISC then activates caspase-8 to be cleaved into its activated form (C8a). Free C8a either forms a complex with a member of the bifunctional apoptotic regulator (BAR) family or triggers the cleavage of Bid into truncated Bid (tBid). From here, tBid promotes mitochondrial permeabilization by activating Bax.^18^ C8a also contributes to apoptosis by catalyzing the activation of caspase-3 (C3) into its activated form (C3a), a well-known executioner of apoptosis. C3a activity is negatively regulated by the inhibitor of apoptotic protein, XIAP, which binds to C3a and inhibits its apoptotic activity by preventing C3a-mediated substrate cleavage (**Figure 2**).

**Figure 2:**
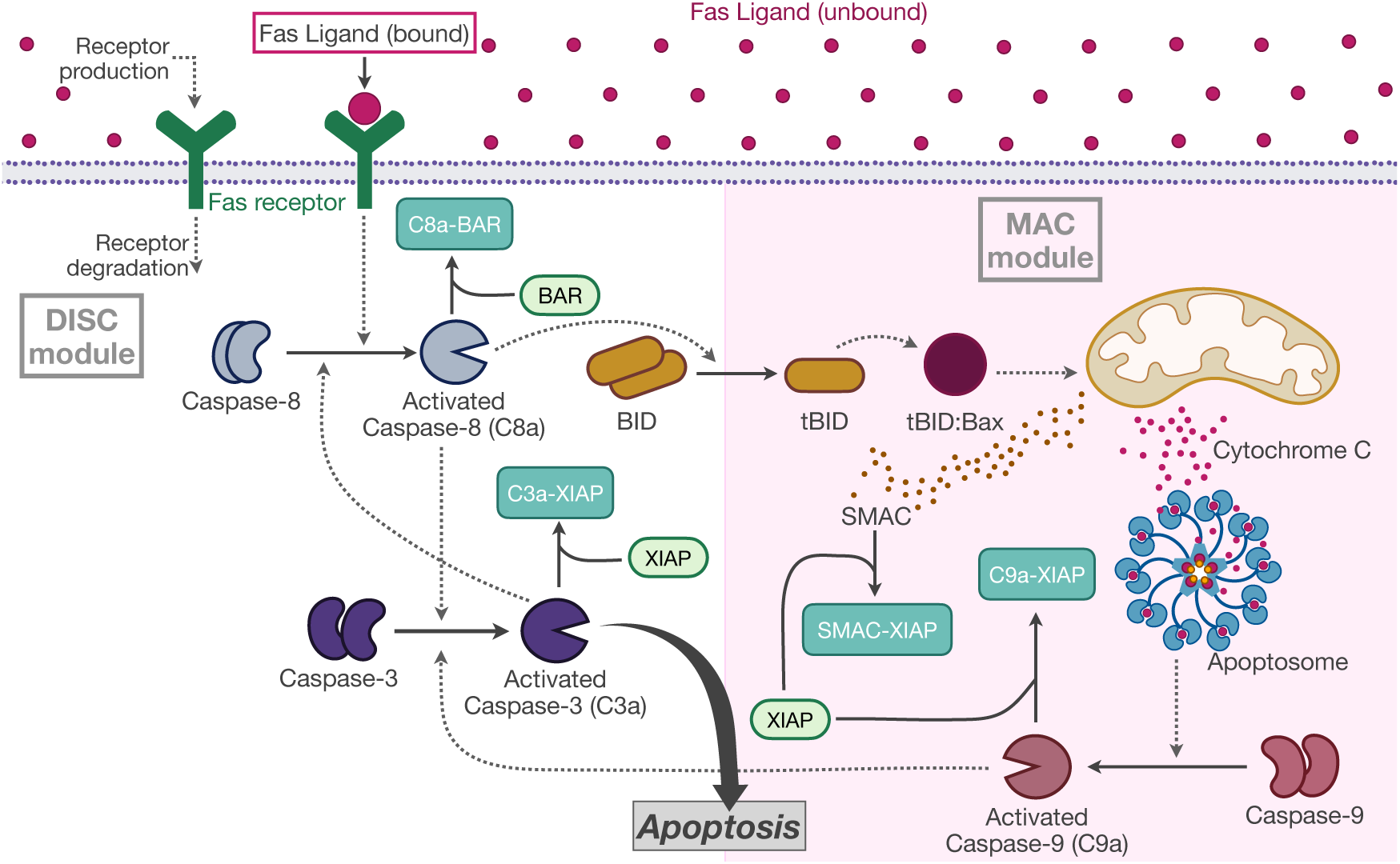
Schematic of the caspase-mediated apoptosis model. The model includes extrinsic and intrinsic (highlighted in pink) caspase-mediated apoptosis model, including the death inducing signaling complex (DISC) and mitochondrial activated caspases (MAC) modules. The binding of FasL to Fas receptor, within the DISC module, creates a complex that then triggers the intracellular pathway by combining with caspase-8 to form activated caspase-8 (C8a). C8a can either combine with caspase-3 to trigger activated caspase 3 (C3a), form a complex made of activated caspase-8 and bifunctional apoptosis regulator (BAR), or initiate the MAC module. In the MAC module, the mitochondrial outer membrane becomes permeabilized, and Cytochrome c (CytoC) and SMAC are released to activate the apoptosome. This leads to further activation of C3a via activated caspase-9. C3a can either trigger apoptosis or form a complex made of C3a and inhibitor of apoptotic protein (XIAP). Solid lines denote protein-protein interactions (e.g., complex formation or binding). Dashed lines denote catalytic interactions, such as enzymatic cleavage or activation of downstream substrates.

Activated caspase-9 (C9a) is the key output of the intrinsic caspase-mediated apoptosis pathway (shaded in pink). C9a promotes activation of caspase-3. XIAP suppresses apoptosis by binding to and inhibiting the activity of C9a. Following activation of tBid, the tBid-Bax axis promotes mitochondrial outer membrane permeabilization, resulting in the release of both Cytochrome c (CytoC) and second mitochondria-derived activator of caspases (SMAC) from the mitochondria. CytoC activates the apoptosome to promote apoptosis. SMAC functions as an antagonist of XIAP by binding and neutralizing XIAP, thereby relieving XIAP-mediated inhibition of caspases and promoting apoptosis. Although there are additional BH3-only proteins besides Bid that affect apoptosis mediated by the intrinsic mitochondrial pathway, we prioritized the Bid/tBid axis since Bid is considered a critical BH3-only protein that links caspase-8 activation to the intrinsic pathway. Additionally, we omit proteins from the Bcl-2 family due to a lack of experimental data needed to specify model parameters.

The model also includes positive feedback between caspase-3 and caspase-8 and -9, based on literature evidence. Prior modeling work of Harrington and coworkers^16^ includes C3a-mediated activation of caspase-8 and caspase-9; therefore, we included this in our model as well. This positive feedback is also described in experimental studies,^19–21^ including a recent investigation of the role of caspase-3 using CRISPR-based gene editing.^22^

For a full list of reactions, ODEs, parameter values, initial conditions, and how each drug was affixed to the model, refer to *Supplemental Tables 1-4*. The model files are available via GitHub: https://github.com/FinleyLabUSC/Apoptosis-drug-resistance.

### Experimental Data

The experimental datasets extracted for model calibration were selected based on the following criteria: experiments were conducted on the same prostate cancer cell line (PC3), data for caspase signaling species were available, and each species had at least three timepoints beginning at 0 hours. Three independent datasets were obtained that met these criteria, where caspase signaling was measured following administration of a pro-apoptotic drug:

● Narciclasine: C8a was measured at 0, 16, 24, and 48 hours, and procaspase-3 (C3) at 0, 16, and 24 hours. Narciclasine was given at a concentration of 1 µM.^23^ Since the authors only observed changes in the levels of C3 (and not C3a), we compare our model simulations for C3, rather than the activated form of caspase-3.
● *Tocopheryloxybutyrate:* C3a and C9a were measured at 0, 24, and 48 hours. Normalized ratios were provided in the publication, so ImageJ analysis was not required. The αTS drug analog was used, given at a concentration of 40 µM.^24^
● *Celecoxib:* C3a was measured at 0, 12, 24, and 48 hours. Celecoxib was given at a concentration of 100 µM.^25^

For each drug, the modeled effects were implemented according to the mechanisms described in the source literature from which the Western blot data were obtained. Narciclasine was modeled as a regulator of upstream extrinsic apoptotic signaling by enhancing the availability of initiator caspases and maintaining them in their active form, based on experimental observations.^23^ In the model, C8a is reversibly sequestered by the inhibitory protein BAR, forming an anti-apoptotic C8a-BAR complex (Supplemental Table 1; *see Reaction 6*). This interaction limits the amount of free C8a available for downstream signaling of C3. To represent this modality for Narciclasine, we included a parameter (*k*_Narc_) to enhance the reverse reaction rate regulating release of C8a from the inhibitory complex. Under this condition, Narciclasine accelerates dissociation of the inhibitory C8a-BAR complex, thereby increasing the pool of free C8a available to activate C3.

Celecoxib is modeled as a pro-apoptotic drug through enhanced removal of inhibitory proteins. Based on evidence that celecoxib can downregulate XIAP, thereby facilitating caspase activation^25^, the model implements Celecoxib as a multiplicative factor (*k*_Cele_) applied to all reactions that reduce XIAP levels in its respective ODE. By scaling all negative terms in the ODE for XIAP removal terms by a value *k*_Cele_ > 1, Celecoxib effectively reduces the pool of inhibitory proteins, increasing the availability of free C3a and promoting apoptotic signaling. This formulation captures the context-dependent pro-apoptotic effect of Celecoxib within the network, allowing it to mitigate XIAP-mediated suppression.

While Narciclasine and Celecoxib promote apoptosis by mitigating the effects of the apoptosis inhibitors BAR and XIAP, respectively, Tocopheryloxybutyrate promotes apoptosis through upregulation of caspase-3.^3^ However, among the three drugs applied to the model, Tocopheryloxybutyrate differed fundamentally from Narciclasine and Celecoxib in how it was simulated. The experimental data revealed that Tocopheryloxybutyrate produced a qualitatively distinct dynamic response. Specifically, the caspase-3 activation profile following Tocopheryloxybutyrate treatment exhibited a transient, bell-shaped response from time 0 hours to 48 hours. In contrast, untreated simulations displayed delayed activation followed by a rapid rise and sustained plateau. This fundamental difference in temporal behavior could not be reproduced by parameter modulation alone. As a result, Tocopheryloxybutyrate required the introduction of an entirely new time-decaying drug input term within the ODE for C3a:

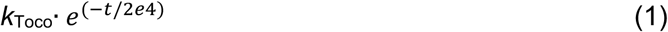

where *k*_Toco_, the initial drug magnitude, is multiplied by an exponential decay term to model loss of efficacy over time.

To reproduce the experimentally observed bell-shaped dynamics of C3a following treatment, two more modifications were introduced: first, *k*_Toco_ was added to the activation of C3a by activated caspase-8 in reaction 3; second, a multiplicative sink parameter, *β*, was included to scale the C3a removal rate. These adjustments allow C3a production to dominate initially, before eventually falling below the clearance rate (via complex degradation), generating the characteristic transient peak.

Mechanistically, this implementation of the effect of Tocopheryloxybutyrate produces three sequential phases: initial accumulation due to production, a slowing and eventual peak as the sink term becomes significant, and a decline as irreversible removal exceeds production. The magnitude of *β*, determines the timing and sharpness of the transient response: lower values produce broad or plateaued peaks, whereas higher values prevent accumulation to C3a levels that surpass untreated cells. This combination of drug-driven production and nonlinear, C3a-dependent, removal successfully generates a bell-shaped C3a profile in qualitative agreement with experimental observations (Supplemental Figure 1).

### Image J

ImageJ is a free and open source programming software that processes scientific images for experimental analysis.^26^ In this work, ImageJ was used to extract the relative amounts of caspase proteins treated with one of three pro-apoptotic prostate cancer drugs from Western blot assay images.^23–25^ Pixel intensity within the region of interest was quantified at each time point by first calibrating the image scale using a blank area of the assay. The data provided in the published papers are given as relative band intensities, rather than absolute concentrations. We elected to normalize the intensities obtained from ImageJ to the final time point, to be consistent across datasets. The output of this approach is a set of quantitative values for the relative levels of specific molecular species involved in apoptosis signaling measured in PC3 cells across several time intervals.

### Extended Fourier Amplitude Sensitivity Testing

Extended Fourier Amplitude Sensitivity Testing (eFAST) is a global sensitivity analysis method used to rank model inputs according to their influence on specific outputs of interest.^27^ This approach is widely applied in biological systems modeling and ODE-based simulations because it quantifies how uncertainty in model inputs propagates to variability in model predictions. In eFAST, each model parameter, defined here as a kinetic rate or a non-zero initial protein concentration, is assigned a unique sinusoidal sampling frequency. In addition, a dummy variable is included, which serves as a baseline measure of numerical noise and sampling uncertainty. Inputs are sampled across user-defined parameter ranges and model outputs are analyzed using Fourier decomposition to attribute output variance to individual input frequencies.

Through this variance decomposition framework, eFAST produces two key sensitivity metrics: the first-order sensitivity index (Si) and the total-order sensitivity index (STotal). The first-order sensitivity, Si, quantifies the fraction of output variance that can be explained solely by variation in a single input parameter, reflecting its direct effect on the model output. In contrast, the total-order sensitivity, STotal, captures both the direct contribution of a parameter and its higher-order interaction effects with all other model inputs. Sensitivity indices are evaluated relative to the dummy variable and, therefore, parameters with indices exceeding that of the dummy variable are considered to have statistically meaningful influence on the model’s behavior.

### Particle Swarm Optimization

Particle swarm optimization (PSO) is a parameter estimation method.^28^ The approach begins with a population of candidate parameter sets, referred to as particles. As these particles move through the parameter space, the objective function is evaluated at each location, and information is shared among particles to identify those yielding the lowest objective function values. Inspired by the coordinated movement of a flock of birds searching for food, PSO iteratively guides the population toward an optimal solution.

In this study, we use the algorithm designed by Iadevaia and coworkers.^29^ The sum of squared error (SSE; Eq. (2)) between model simulations and experimental data was used as the objective function:

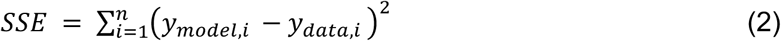

where *n* is the number of experimental datapoints, *y_model_*_,*i*_ is the model-predicted value at the *i*th data point and *y_data_*_,*i*_ is the corresponding experimentally measured value. In addition, the upper and lower bounds are specified for each model input to constrain the possible values the input can take on. PSO was applied to estimate model parameter values that enable the model to capture how pro-apoptotic drugs modulate apoptotic signaling. Only parameters identified as highly influential based on eFAST were estimated using PSO, while all other parameters were fixed to values from the shared baseline model.

The PSO algorithm was run to obtain 100 values for each parameter to ensure robust parameter estimation. Because PSO is a stochastic optimization algorithm, multiple runs help avoid parameter sets from converging at local minima by initializing the particle positions in multiple randomly selected positions. Yet, it is still possible for multiple randomly selected positions to converge to the same observed output. The repeated identification of the same global optimum across many PSO runs demonstrates the stability and reliability of the inferred drug-specific parameter sets. Thus, the final parameter sets were selected based on the runs that achieved the lowest objective function fitting error. It is important to note that using PSO, we simultaneously fit the datasets for the three drugs to obtain estimated model parameters that allow the model to match all of the experimental data.

### Temporal Scaling of the Harrington Model for Prostate Cancer Dynamics

When first running the ODE model of intrinsic and extrinsic apoptosis pathways, the dynamics of the caspases was revealed to be significantly faster than the data from PC3 cells. Specifically, the experimental data showed that caspase activation occurred at approximately 24 hours following drug treatment, while the model simulations showed the activation occurred on the order of minutes. To match this timing, we scaled all kinetic parameters taken from the model by Harrington and coworkers to align with the observed temporal dynamics for the PC3 prostate cancer cell line. Supplemental Figure 2 shows the baseline protein dynamics overlaid with our scaled version, utilizing a multiplicative factor of 0.02 for the kinetic parameters, which produced an appropriate caspase activation timing.

Additionally, we observed strong coupling between baseline initial protein amounts (particularly C3, C8, and C9) taken from the work of Harrington and coworkers and the corresponding protein synthesis rates. To address this, synthesis rate parameters were scaled differently to avoid multiple parameter sets from producing similar outputs, indicating non-identifiable solutions and distorted caspase dynamics. To decouple initial conditions from synthesis-driven accumulation, synthesis rates were scaled to 0.2 rather than 0.02.

### Characterizing apoptotic efficacy

C3a was chosen as the primary actuator of apoptosis because extensive experimental and computational studies indicate that both the intrinsic and extrinsic apoptotic pathways converge at caspase-3 to irreversibly commit the cell to death.^30^ C3a therefore provides a robust and integrative readout of apoptotic activation across all three upstream signaling modules. C9a was selected as a secondary marker of apoptosis given its role as the intrinsic pathway’s gatekeeper to apoptosis as even small variations in C9a activation have been shown to produce large differences in apoptotic activity.^31^

## Data Availability

The data generated in this study were produced using the code publicly available in Github at https://github.com/FinleyLabUSC/Apoptosis-drug-resistance.

## Results

We first examined the key drivers of the baseline model (**Figure 2**), which represents the apoptotic system in the absence of drug treatment (hereafter referred to as the untreated model). This allows us to address the central aims of this study: to evaluate whether the model reliably reproduces apoptosis dynamics under three pro-apoptotic prostate cancer drugs and identify pathway elements that may serve as targets for overcoming treatment resistance.

### Global Sensitivity Analysis Reveals Key Regulators of Apoptosis Signaling

To understand which model inputs (kinetic parameters and initial protein amounts) were most influential in governing C3, C3a, C8a, and C9a outputs across five discrete time points consistent with those used in the experimental data, we performed sensitivity analysis using eFAST on the untreated model first. With all drugs turned off, we varied the kinetic parameters four orders below the baseline and two orders of magnitude above the untreated model while the initial protein amounts were varied two orders of magnitude below and above their respective baseline. This analysis identifies which parameter and initial protein sensitivities most strongly influence the model outputs (**Figure 3**).

**Figure 3:**
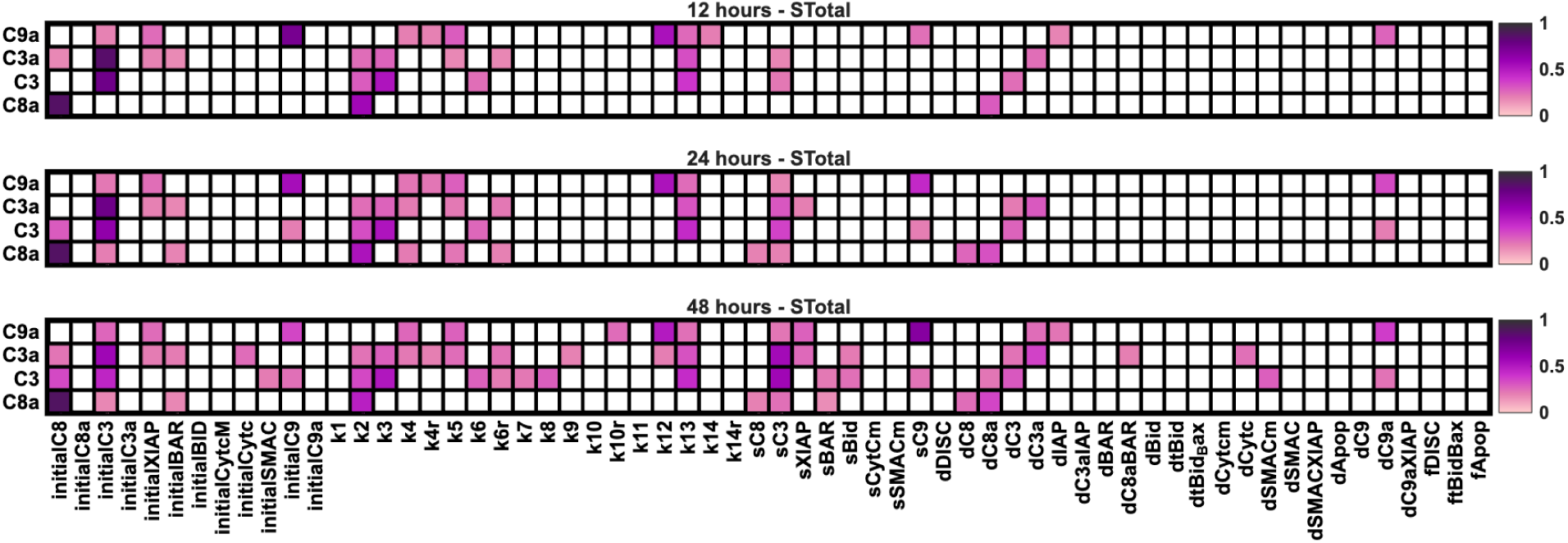
Sensitivity analysis for the untreated model. Heatmaps depict the total sensitivity (STotal) indices for initial concentrations and kinetic parameters in the untreated model, listed along the horizontal axis, for different simulated times. Outputs are caspases, listed along the vertical axis. Lighter pink shades correspond to lower sensitivity values, whereas darker purple shades indicate higher sensitivity. White regions represent parameters whose STotal is not statistically significant when compared against the dummy variable.

To rank the sensitivity indices, we considered STotal thresholds from 0.4 to 0.8 to quantify which parameters and initial conditions were the main drivers across the entire simulation time. For the untreated model, inputs exhibiting STotal ≥ 0.40 included initial C8, initial C3, the rate of C8 activation by C3a (k2), the rate of C3 activation by C8a (k3), the rate of C9 activation by C3a (k12), the activation of C3 by C9a (k13), the synthesis rate of C3 (sC3), the synthesis rate of C9 (sC9), and the degradation rate of C3a (dC3a). At the highest threshold examined (STotal ≥ 0.80), outputs were almost entirely driven by initial C8 in the untreated model.

Next, we performed eFAST with each drug applied individually to identify which model inputs became most influential when the system was perturbed by a specific treatment. (**Figure 4**) When Tocopheryloxybutyrate was active, the STotal ≥ 0.40 parameters included the initial amounts of C8 and C3, k2, k3, k12, k13, sC3, and sC9. Unlike the untreated model, dC3a was also a member of this subset for Tocopheryloxybutyrate. Celecoxib exhibited a largely overlapping sensitivity profile, with the same influential parameters as Tocopheryloxybutyrate except for the degradation of C3a and k13, whose STotal did not exceed 0.40 under Celecoxib treatment. When Narciclasine was active, model sensitivity was influenced by the same threshold by a subset of seven parameters, including the initial amounts of C8 and C3, k2, k12, k13, sC3, sC9.

**Figure 4:**
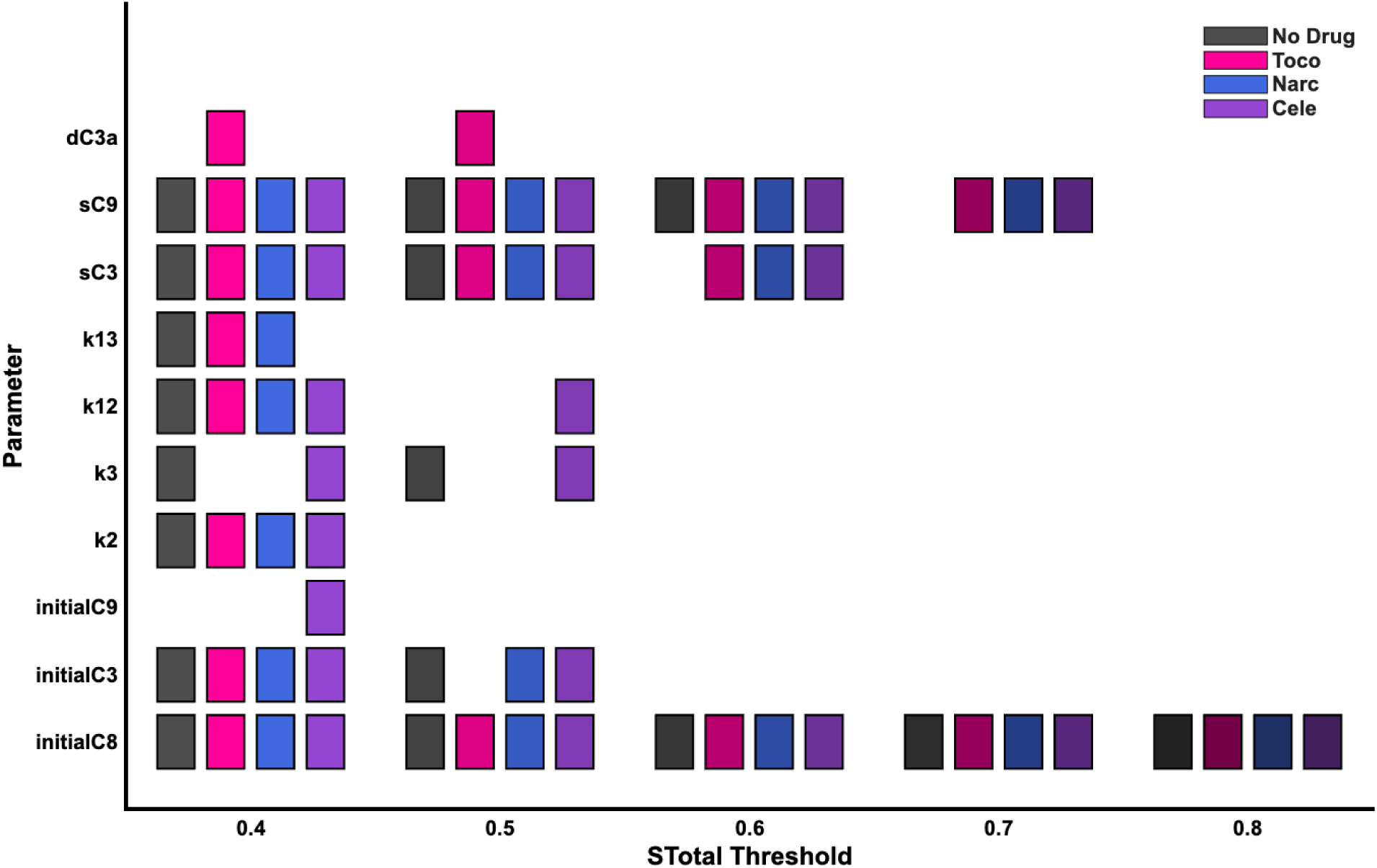
Total sensitivity indices. Identification of influential parameters, those with a STotal sensitivity index exceeding a given threshold ranging from 0.4 to 0.8. Stotal values are shown for the untreated case (black) or with Tocopheryloxybutyrate (pink), Narciclasine (blue), or Celecoxib (purple) treatment. Darker shading indicates increasing STotal threshold values.

As a supplemental analysis, we compared first-order and total-order sensitivity indices demonstrating the importance of parameter interactions within the system. While initial amounts of C3 consistently exhibited a high first-order sensitivity index across all thresholds, more model inputs satisfied the total-order index threshold (Supplemental Figure 3). This disparity indicates that apoptotic signaling behavior is strongly influenced by nonlinear interactions between kinetic parameters and initial protein amounts, rather than isolated model inputs.

### The Model Captures the Qualitative Dynamics of Three Pro-Apoptotic Drugs

The eFAST analysis identified 10 parameters and initial conditions that met the STotal ≥ 0.4 threshold for each of the four treatment conditions (**Figure 4**): initial C8, initial C3, initial C9, k2, k3, k12, k13, sC3, sC9, dC3a. We next aimed to identify the optimal values for these 10 model inputs using PSO, simultaneously fitting the model to all three pro-apoptotic experimental datasets with each of the drugs turned on. Notably, the bounds for each input were kept consistent with the ranges defined during eFAST. Experimental measurements for the caspase levels at each time point were quantified using ImageJ. The SSE value for each PSO run can be found in Supplemental Figure 4. The best-fit parameter sets were extracted from the 100 runs to include runs that did not contain any parameters estimated to be at their respective lower or upper bound (Supplemental Figure Figure 5), resulting in 60 parameter sets. The estimated parameter values are given in Supplemental Table S5. We simulated the caspase dynamics using the 60 best-fit parameter sets, and we show the mean of the model simulations over time. The calibrated model quantitatively and qualitatively the data from the three datasets (**Figure 5**), the relative levels of the signaling species over time. With the calibrated model, we can investigate the behavior for a range of conditions, including varying drug strengths and with combination treatment. Furthermore, we explore how the initial protein amounts correspond to the level of C3 activation.

**Figure 5:**
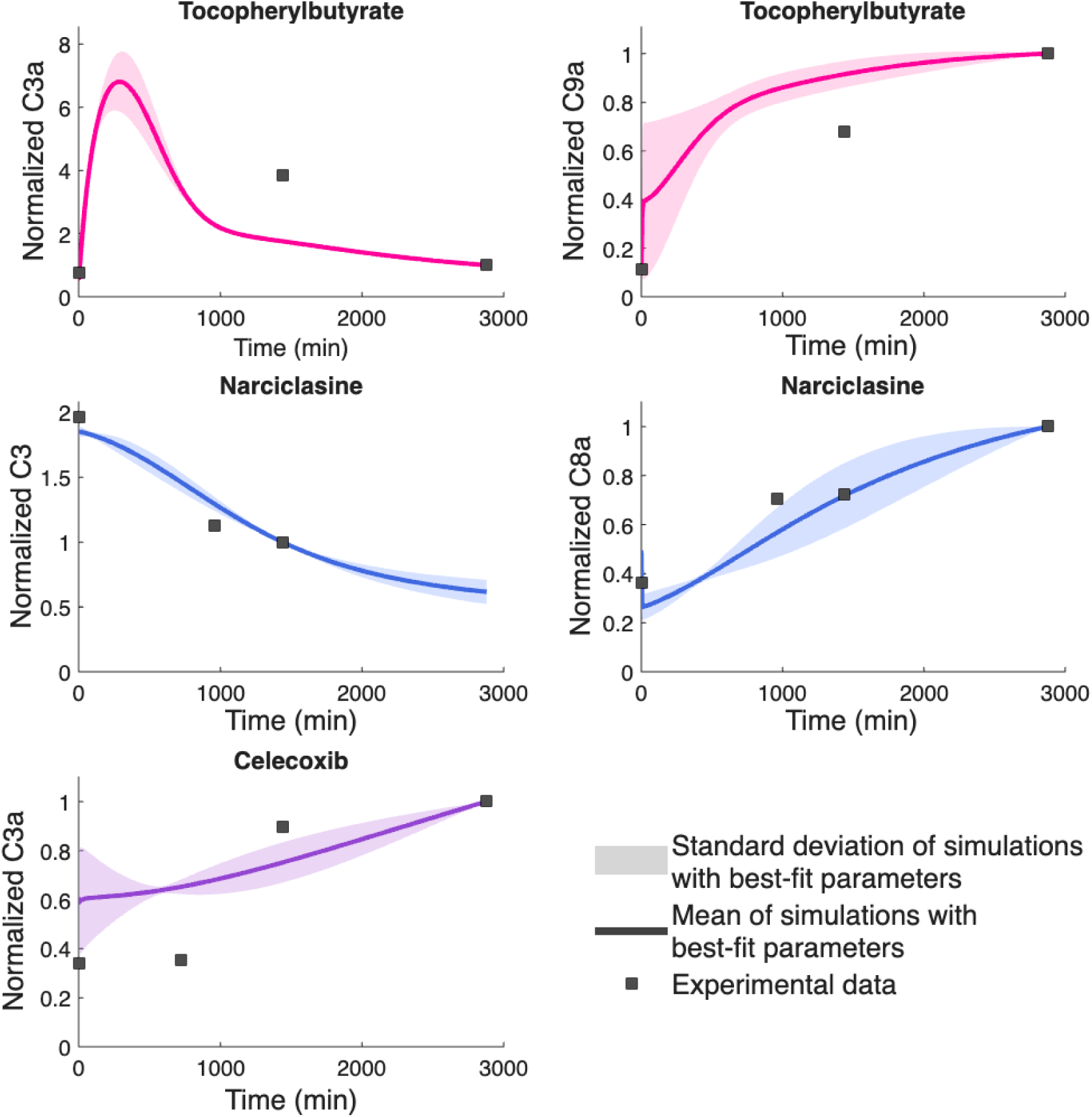
Model simulations using the parameters estimated with PSO. Experimental data for C3, C3a, C8a, and C9a were simultaneously fit to experimental data (black squares) quantified using ImageJ. Tocopheryloxybutyrate data include C3a and C9a measured at 0, 1440, and 2880 minutes. Narciclasine data include C3 measured at 0, 960, 1440, and 2880 minutes and C8a measured at 0, 960, and 1440 minutes. Celecoxib data include C3a measured at 0, 720, 1440, and 2880 minutes. Solid line is the mean value of the model simulations. Shaded regions represent the standard deviation across the best-fit parameter sets.

To demonstrate that the model is not limited to a single treatment or cell line, we compared model predictions for caspase dynamics to experimental data from a separate prostate cancer cell line and a different analog to the Tocopheryloxybutyrate drug.^23^ These data were obtained from the LNCaP cell line for the αTS drug analog (the same as used for model fitting above) and the αTOB drug analog, along with the αTOB drug analog in the PC3 cell line. Supplemental Figure 6 highlights the qualitative agreement between model predictions and experimental data across cell lines. Furthermore, these results support future integration with cell-specific omics data to refine parameterization, further enhancing the model’s predictive capacity across cell lines and drug targets.

Overall, the PSO algorithm produces a set of parameters that simultaneously matches the available experimental data used for fitting (**Figure 5**) and qualitatively matches data not included for fitting (Supplemental Figure 6). Thus, we have a calibrated model that gives reliable simulation results. We next applied the calibrated model to simulate conditions that have not been studied experimentally and for which experimental data are not available: varying the strength of the drugs, simulating drug combinations, and evaluating the effect of varying the signaling species’ initial concentrations. Results from these model simulations are presented below.

### Effector Caspase Activations Increase as Drug Strength Increases

Using the 60 best-fit parameter sets identified by PSO, we used the calibrated model to examine how drug strength influences the apoptotic effector, C3a. We simulated the model we varied the strength of the three anti-apoptotic drugs (**Figure 6**). For Tocopheryloxybutyrate (*k*_Tele_ ranging from 1×10⁻³ to 9×10⁻³), increasing the drug strength led to the highest fold-change in C3a, compared to the other drugs. For varying strength of Narciclasine (*k*_Narc_ ranging from 1 to 9), higher drug strengths induced more activation of C3a. In contrast, changing the strength of Celecoxib (*k*_Cele_ varying from 1 to 9) produced minimal effects on the apoptotic markers. Overall, drug strength was positively correlated with C3a levels, with higher drug intensities producing greater activation of the apoptotic actuators for each pro-apoptotic drug.

**Figure 6:**
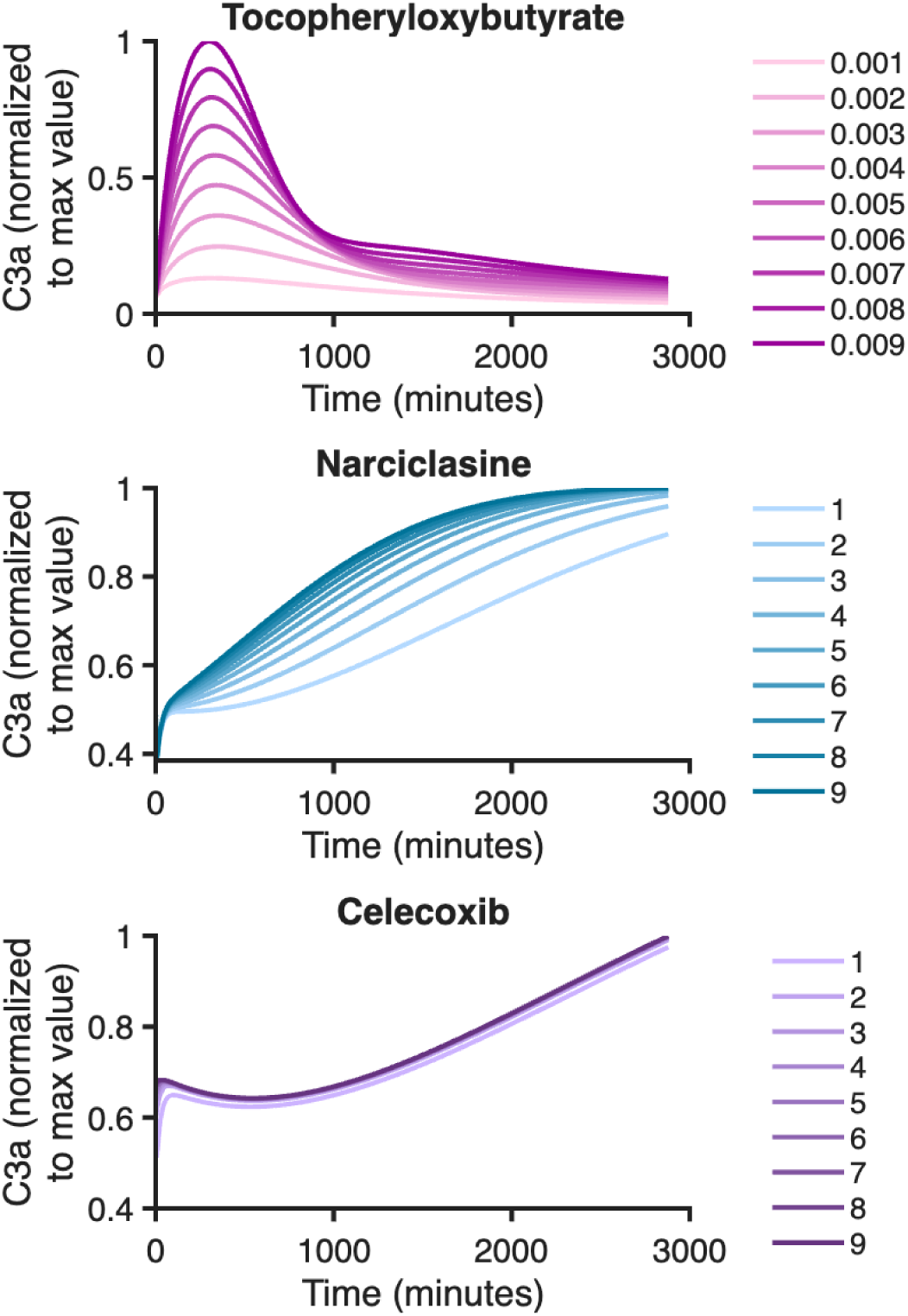
Effects of varying drug strength on effector caspase dynamics. Activated caspase-3 (C3a) is shown, with varying strength of a single drug, with the other two drugs turned off. Line color indicates drug strength. C3a values have been normalized to the maximum C3a response induced by the highest drug strength simulated, for visualization purposes.

### Combination Drug Therapy Reveals Differential and Nonlinear Effects on C3a

Once the effects of individual drug strength were characterized, we next evaluated whether apoptotic activation could be enhanced through combination therapy. Because C3a was operationally defined as the model’s downstream actuator of apoptosis, peak C3a levels were quantified across the same range of drug strengths described above for all pairwise and triple-drug combinations. Furthermore, we normalized to the peak C3a from the untreated model.

Combination treatments demonstrated pronounced, non-additive C3a levels, compared to no-treatment (**Figure 7**). Combined treatment with Tocopheryloxybutyrate and Narciclasine slightly reduced the relative maximal C3a response compared to Tocopheryloxybutyrate alone (max is 6.09×10⁵ compared to 8.01×10⁵), suggesting an antagonistic or competitive interaction within the apoptotic signaling network. In contrast, Tocopheryloxybutyrate combined with Celecoxib produced a maximal response (8.06×10⁵) slightly exceeding the effect of Tocopheryloxybutyrate alone, indicating minimal interference between the drugs. The Narciclasine and Celecoxib pairing remained confined to the low-amplitude regime, similar to Narciclasine implemented alone. Notably, the triple-drug combination implementation closely mirrored the Tocopheryloxybutyrate and Narciclasine response (max relative C3a is 6.09×10⁵), further supporting the presence of inhibitory modulation associated with Narciclasine under certain conditions.

**Figure 7:**
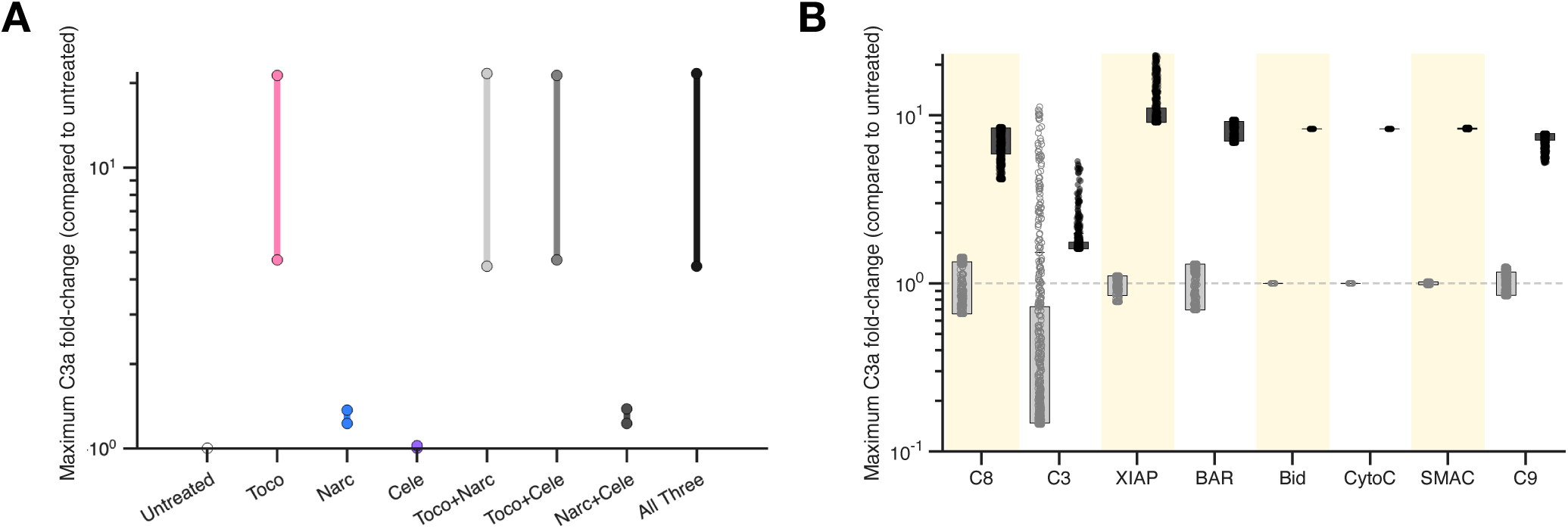
Effect of drug treatment considering cell heterogeneity. (**A**) Comparison of the range of maximum C3a response produced by the calibrated model for single and combination drug treatments. Maximum C3a is normalized to the untreated condition. The vertical bars represent the full range of the maximum C3a values observed across all drug strengths. (**B**) The maximum C3a responses across the range of initial protein levels for 500 simulated cells, relative to mean of the untreated condition. The responses for the triple-drug combination (dark grey) are compared to the untreated condition (light grey open circles).

### Intracellular Protein Heterogeneity Predicates Therapeutic Efficacy

Following the observation that combination therapies exhibited non-additive and, in some cases, attenuated responses, we next investigated how variability in intracellular protein amounts influences therapeutic efficacy. To this end, we quantified the intrinsic influence of key apoptotic protein amounts on maximum C3a activation across two conditions: the untreated model (all drugs inactive) and the combination therapy model (all three pro-apoptotic drugs active), with intracellular protein levels varied across simulated cell populations.

We simulated 500 heterogeneous cells, each having different initial protein amounts. Specifically, we individually varied the amount of C8, C3, XIAP, BAR, Bid, CytoC, SMAC, and C9 by sampling over a log-uniform range spanning two orders of magnitude around the baseline values taken from the model by Harrington and coworkers. Besides the individual protein whose initial amount was being varied, all other initial conditions and model parameters were set to the PSO-derived best-fit value with the lowest SSE. Maximum C3a levels were computed for both conditions and normalized to the mean no-drug baseline. Boxplots represent the distribution of fold-change responses across simulated cells, with the dashed line indicating the untreated condition (fold-change = 1). When considering the initial pro-caspase levels, there is no overlap in initial C8 or C9 levels between combination therapy and untreated conditions (**Figure 9**). In contrast, initial C3 levels do not exhibit such a distinct disparity between the two conditions as some simulations from both groups have shared C3a amounts. Additionally, distinct initial conditions for Bid, CytoC, and SMAC correspond to untreated and combination treatment, where higher initial concentrations of these proteins are associated with increased drug-induced C3a activation. This result is intuitive, as these proteins do not inhibit caspase-3 activation. Interestingly, higher initial levels of the inhibitors of apoptosis signaling, BAR and XIAP also correspond to higher C3a activation with treatment.

## XIAP Levels Constrain Apoptotic Potential

Given the observation that higher initial BAR and XIAP levels were associated with increased drug-induced C3a activation (**Figure 7**), we further examined how variability in these inhibitory proteins impacts apoptotic signaling. We simulated a heterogeneous population of 2,000 cells where each simulation represents a single cell with all non-zero parent proteins independently sampled over four orders of magnitude. As expected, combination therapy increases the maximum C3a concentration, compared to the no-drug baseline (Supplemental Figure 7). We then considered the relationship between initial C3 and the resulting maximum C3a level (Figure 10A). Model simulations show a range of C3a activation for each simulated cell, given varying initial C3 levels. Interestingly, the maximal C3a reached is independent of initial C3 levels below ∼100 nM. Above that initial C3 level, maximal C3a exhibits a strong positive correlation with the initial C3 concentration.

We then considered how the initial levels of XIAP and BAR affect the maximal C3a attained. For the heterogeneous population of cells, the initial C3 level influences maximal C3a more strongly than XIAP (**Figure 8B**, top panel). When considering the combined influence of initial XIAP and BAR levels, in general, a combination of low XIAP and BAR correspond to higher maximal C3a (dark points in **Figure 8B**, bottom panel). However, some combinations of high initial XIAP and intermediate initial levels of BAR also give relatively high maximal C3a. Thus, we further investigated this relationship between the inhibitors of apoptosis and C3a response. We considered the set of simulated cells in which the maximum C3a level exceeded a specified threshold. We selected the threshold to be 200 nM to capture the cells that exhibit a large response following combination therapy (dashed line in **Figure 8A**). Approximately 15% of the simulated cells had a maximum C3a level that met this threshold (291 out of 2,000). These cells had a lower initial XIAP concentration compared to cells in which the maximum C3a level was below this threshold (**Figure 8C**). We observed the same trend for the initial BAR concentration (Supplemental Figure 8). Thus, cells that exhibit a low response to combination therapy have higher initial XIAP and BAR concentrations. This indicates that enhancing effector caspase amounts alone is insufficient to increase apoptosis unless XIAP inhibition is also high. Targeting XIAP is particularly important to shift the cells’ C3a response above a baseline level. Altogether, this analysis identifies that the relative amounts of BAR and XIAP are key regulatory features of the integrated apoptosis pathway, suggesting that modulating these inhibitory proteins could serve as a viable strategy for overcoming treatment resistance.

**Figure 8:**
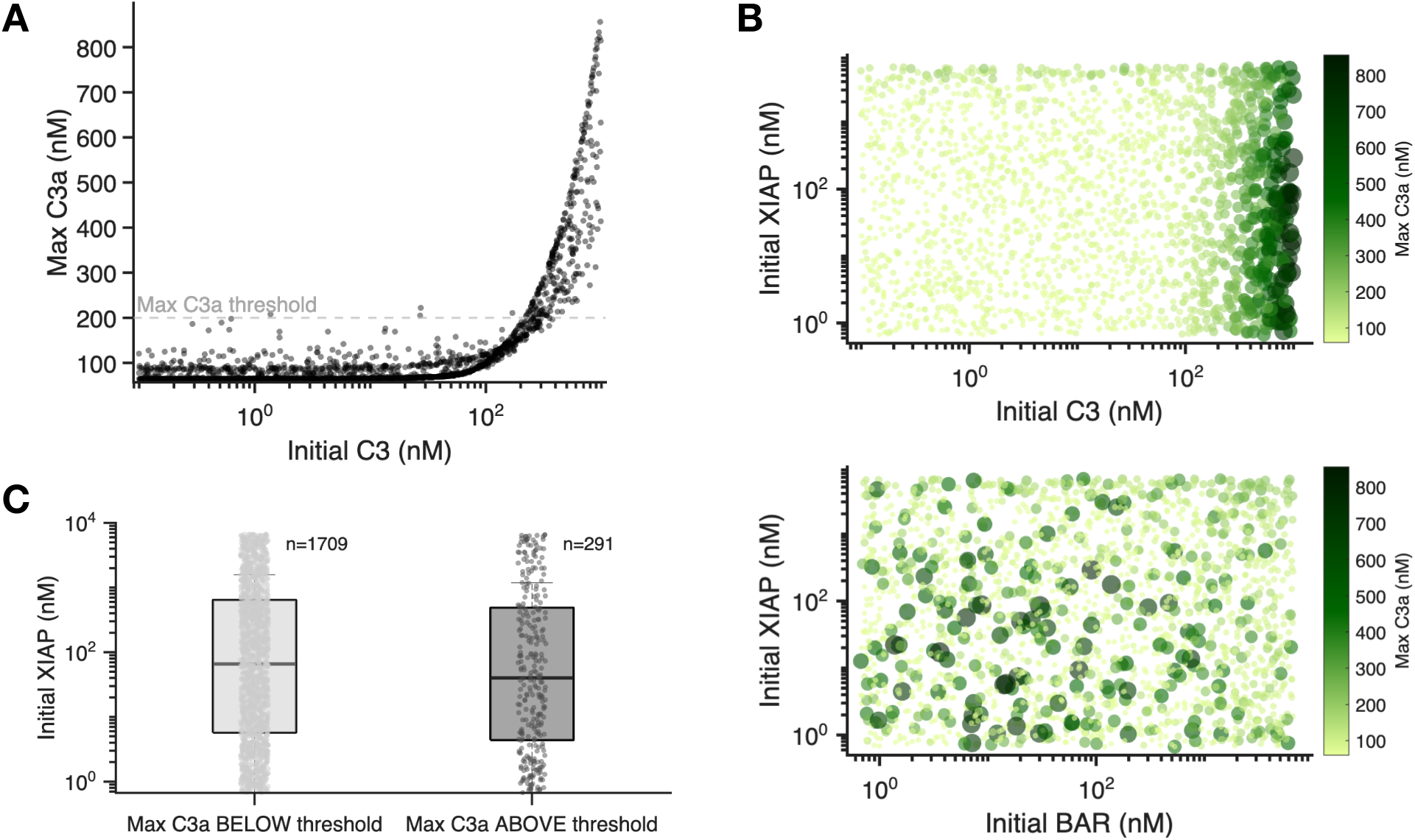
Influence of XIAP and BAR on C3a activation for combination therapy. (**A**) Scatter plot of maximum C3a versus initial C3 across 2000 simulated cells with combination therapy. The dashed line denotes the threshold used for further analysis. (**B**) Scatter plot of maximum C3a as a function of initial C3 and XIAP (top) or initial XIAP and BAR (bottom). Color intensity and dot size represent maximum C3a level. (**C**) Initial XIAP level for cells in which the maximum C3a is below (light grey; median = 65.9 nM) or above (dark grey; median = 39.9 nM) the specified threshold shown in panel A of 200 nM.

## Discussion

For pro-apoptotic therapies to be effective in prostate cancer, apoptotic signaling pathways must remain functional, and pro-apoptotic proteins must not be counteracted by excessive levels of inhibitors of apoptosis (pro-survival proteins). Failure to meet either of these conditions can limit drug efficacy and contribute to intrinsic resistance.^32^ In this work, we demonstrate this by answering two main questions: *(1) Can caspase-mediated apoptosis dynamics be accurately captured in response to three separate prostate cancer pro-apoptotic drugs? (2) Which features of the caspase-mediated apoptosis pathway represent viable targets for overcoming treatment resistance?*

The model designed in this work closely recapitulates the diverse qualitative and drug-specific dynamics across three pro-apoptotic drugs in a prostate cancer cell line, while revealing how drug strength, combination therapy, and the landscape of inhibitory proteins can impact the magnitude of key apoptotic actuators. Leveraging this model, we were able to demonstrate how the apoptotic response to pro-apoptotic drugs is governed by an interplay between kinetic rates, intracellular protein quantities, and the context-dependent state of prostate cancer cells, collectively shaping intrinsic drug resistance. Interestingly, although XIAP emerged as an influential regulatory node in our analysis, its therapeutic targeting has not been shown to effectively promote apoptosis as strongly as expected.^33^ The limited effect of targeting XIAP may be due to compensatory survival mechanisms and pathway redundancy. Additionally, the influence of XIAP is predicted to be context-dependent, where other signaling species contribute to its effect. This highlights the importance of using a systems-level approach to identify effective pro-apoptotic combinations. Excitingly, our model provides a framework to enable such studies.

For instance, Narciclasine exerts its effect by promoting the dissociation of the C8a-BAR inhibitory complex, freeing active C8a to initiate downstream caspase activation in our model. The efficacy of Narciclasine is, therefore, intrinsically linked to the abundance of BAR at the time it is implemented since higher initial BAR levels provide more substrate for complex dissociation, leading to greater release of C8a and subsequently stronger activation of the apoptotic pathway. Yet, when Narciclasine is combined with Celecoxib or Tocopheryloxybutyrate, the model predicts non-linear interactions that are not intrinsically additive. This suggests that Narciclasine’s activity is strongly affected by the levels of anti-apoptotic or pro-survival proteins, indicating potential intrinsic resistance mechanisms.

As another exemplary case, while Celecoxib promotes apoptosis and enhanced C3a activation by reducing XIAP, combination therapy with other pro-apoptotic drugs can still sustain levels of free BAR. This intrinsic dynamic stems from how the system is designed. When C8a is freed from inhibitory complexes, it promotes the activation of effector caspases downstream, but other inhibitory complexes may still thrive in a way that overwhelmingly promotes survival of prostate cancer cells. Thus, the model quantitatively explains why the effect of a drug is not solely dependent on its direct target. Instead, the drug effect also depends on the cell’s intrinsic starting state and the abundance of other promotors and inhibitors of apoptosis. This phenomenon highlights why it is important to consider cellular context and drug combinations when exploring apoptosis signaling and designing effective therapies.

Consequently, because apoptotic regulation and therapeutic response are governed by a complex network of interacting factors, computational modeling has emerged as a powerful tool to systematically investigate these effects. For example, computational models have guided the discovery of therapeutic molecules that restore p53, a tumor suppressor gene critical for initiating apoptosis when DNA damage is irreparable.^34^ *In silico* models have elucidated how survival signals can inhibit apoptotic activation and have even shown how psychological stress can disrupt cancer cell death.^35,36^ Building on these applications, the model described here provides a transferable framework that simulates prostate cancer cell apoptosis signaling without relying on hormonal signaling. The model thus enables progress toward optimizing the efficacy of apoptosis-targeting therapies.

Our model integrates the intrinsic and extrinsic caspase-mediated apoptotic pathways, accounting for cross-amplification of apoptosis signaling and enabling design of therapeutic strategies that act on these pathways. Simulated the combined effects of the intrinsic and extrinsic pathways is critical because the two pathways have distinct levels of activation across different cell types. Specifically, cells known as *type II* cells, require amplification through the intrinsic pathway for effective apoptosis, while *type I* cells can complete extrinsic apoptosis without engaging the intrinsic pathway.^37^ Prior computational apoptosis models typically evaluate mechanisms of either the intrinsic or extrinsic pathway alone, as jointly calibrating both pathways requires extensive parameterization and experimental data. Excitingly, in this work we successfully identify influential model parameters and estimate their values by fitting to protein measurements from a prostate cancer cell line (PC3).

However, we acknowledge limitations of this work. First, we made some assumptions in model development, which influence the structure of the model. We aimed to construct and calibrate a parsimonious model of caspase-mediated apoptosis, excluding some species known to influence apoptosis. For example, we account for Bid/tBid rather than other BH3-only proteins such as Bim, Puma, and Noxa. We prioritized the Bid/tBid axis given that Bid is considered a critical BH3-only protein connecting caspase-8 activation to the intrinsic mitochondrial pathway that affects apoptosis.^38,39^ Similarly, we do not include the anti-apoptotic Bcl-2 family proteins that influence the mitochondrial pathway. As the present model was designed to capture the core signaling events supported by available experimental datasets in prostate cancer cell lines, model simulations are limited by the availability of experimental data. Including additional signaling species would increase the number of model parameters whose values would need to be estimated using experimental data, which is limited. Thus, we simplified our model to align with data availability. Future work can expand the model to include other BH3-only proteins such as Bim, Puma, and Noxa, as well as proteins from the Bcl-2 family.

A second limitation is that we present a computational model calibrated using data from a single prostate cancer cell line. We address this limitation by evaluating the model’s applicability across additional experimental conditions, using separate data from the LNCaP cell line and a different analog to the Tocopheryloxybutyrate drug (Supplemental Figure 5). The comparisons between model predictions and these independent datasets indicate that the model captures key features of apoptotic regulation beyond the training data used for fitting. Additionally, because the model is grounded in conserved mechanisms of caspase activation, it can be extended to related cellular contexts through parameter re-estimation rather than requiring modifications to the structure of the model itself. The results of this work support the potential of the model as a flexible platform for exploring responses across additional cancer subtypes and treatment conditions.

Building on this foundation, future work aims to improve predictive accuracy. This includes generating additional experimental datasets in the PC3 cell line (untreated and with additional drugs) to better constrain model parameters and capture variability in drug response. In addition to immortalized cell lines, patient-derived tumor organoids are increasingly used to provide a more widely used experimental model for studying drug resistance. Our model can also be fit to data generated from organoid studies, providing a more human-relevant modeling framework. Furthermore, it will be important to use additional cell lines and tumor organoids to validate the model findings, such as the impact of XIAP in combination with other signaling species. Also, incorporating the mitogen activated protein kinase (MAPK) pathway will provide more insight into the specific context of prostate cancer biology and survival signaling crosstalk with apoptosis signaling. Finally, extending and validating the model in other hormone-dependent cell types will allow us to evaluate its broader translatability and determine whether the identified regulatory mechanisms are conserved across systems or specific to prostate cancer cells.

There is an urgent need for treatment strategies that circumvent hormonal signaling pathways in hormone-dependent cancers, as advanced prostate cancer cell lines such as PC3 often progress to an androgen-independent state.^40^ In this study, we have presented an *in silico* model that quantitatively characterizes how suppression of caspase-mediated apoptosis machinery contributes to dysfunctional cell-death signaling, while revealing therapeutic vulnerabilities across kinetic parameters and initial protein amounts. Altogether, our findings emphasize that intrinsic drug resistance is often the result of synergistic and competing interactions between multiple regulatory processes, rather than from the influence of individual components acting in isolation. This phenomenon is not unique to cancer signaling and, importantly, is not limited to prostate cancer cells. The same framework can be extended to other cancer types, especially characteristically incurable cancers, where apoptosis resistance is one of the major mechanisms that allows cancer cells to persist once tumorigenesis begins. Ultimately, such models can noninvasively and rapidly evaluate personalized therapeutic strategies tailored to the unique biological conditions that account for the complex heterogeneous profile of cancer cells. Therefore, this work lays the groundwork for designing targeted therapeutics in a wide range of cancer types.

## Acknowledgements

The authors acknowledge members of the Finley research group for constructive feedback.

## Study Highlights

Apoptosis is a promising therapeutic target in castration-resistant prostate cancer, but the complexity of apoptosis signaling and the emergence of intrinsic and acquired resistance make it difficult to identify the molecular determinants of treatment response. This work answers the question “Can a reduced, data-calibrated mechanistic model of apoptosis signaling identify key regulators of drug response and resistance in prostate cancer?” We developed a parsimonious ordinary differential equation model calibrated to experimental data from three pro-apoptotic agents. The model reproduced observed apoptotic responses, identified critical determinants of apoptotic commitment and resistance, and generated testable hypotheses regarding therapeutic vulnerabilities and effective drug combinations. This framework provides a quantitative platform for investigating resistance mechanisms, prioritizing therapeutic targets, and evaluating combination strategies in silico, thereby supporting more efficient development of apoptosis-targeting therapies and guiding future experimental studies.

## Author Contributions

DSM and SDF wrote manuscript, designed research and analyzed data; DSM performed research.

## References

1. Maldonado, E. B., Parsons, S., Chen, E. Y., Haslam, A. & Prasad, V. Estimation of US patients with cancer who may respond to cytotoxic chemotherapy. Future Sci. OA 6, FSO600 (2020).

2. Glenfield, C. & Innan, H. Gene Duplication and Gene Fusion Are Important Drivers of Tumourigenesis during Cancer Evolution. Genes 12, 1376 (2021).

3. Liu, M. et al. Celecoxib regulates apoptosis and autophagy via the PI3K/Akt signaling pathway in SGC-7901 gastric cancer cells. Int. J. Mol. Med. 33, 1451–1458 (2014).

4. Dhakne, P. et al. Refinement of safety and efficacy of anti-cancer chemotherapeutics by tailoring their site-specific intracellular bioavailability through transporter modulation. Biochim. Biophys. Acta Rev. Cancer 1878, 188906 (2023).

5. Utku, N. New approaches to treat cancer - what they can and cannot do. Biotechnol. Healthc. 8, 25–27 (2011).

6. Oliver Metzig, M. & Hoffmann, A. Controlling Cancer Cell Death Types to Optimize Anti-Tumor Immunity. Biomedicines 10, 974 (2022).

7. Di Maggio, F. M. et al. Portrait of inflammatory response to ionizing radiation treatment. J. Inflamm. 12, 14 (2015).

8. Hanahan, D. Hallmarks of cancer-Then and now, and beyond. Cell 189, 2254–2277 (2026).

9. Ali, A. & Kulik, G. Signaling Pathways That Control Apoptosis in Prostate Cancer. Cancers 13, 937 (2021).

10. Mangrum, D. S. & Finley, S. D. Modeling the heterogeneous apoptotic response of caspase-mediated signaling in tumor cells. J. Theor. Biol. 590, 111857 (2024).

11. McComb, S. et al. Efficient apoptosis requires feedback amplification of upstream apoptotic signals by effector caspase-3 or -7. Sci. Adv. 5, eaau9433 (2019).

12. Li, K., van Delft, M. F. & Dewson, G. Too much death can kill you: inhibiting intrinsic apoptosis to treat disease. EMBO J. 40, e107341 (2021).

13. Key Statistics for Prostate Cancer | Prostate Cancer Facts | American Cancer Society. https://www.cancer.org/cancer/types/prostate-cancer/about/key-statistics.html.

14. Karantanos, T., Corn, P. G. & Thompson, T. C. Prostate cancer progression after androgen deprivation therapy: mechanisms of castrate resistance and novel therapeutic approaches. Oncogene 32, 5501–5511 (2013).

15. Nakazawa, M., Paller, C. & Kyprianou, N. Mechanisms of Therapeutic Resistance in Prostate Cancer. Curr. Oncol. Rep. 19, 13 (2017).

16. Harrington, H. A., Ho, K. L., Ghosh, S. & Tung, K. C. Construction and analysis of a modular model of caspase activation in apoptosis. Theor. Biol. Med. Model. 5, 26 (2008).

17. Hass, H. et al. Benchmark problems for dynamic modeling of intracellular processes. Bioinformatics 35, 3073–3082 (2019).

18. Flores-Romero, H. et al. BCL-2-family protein tBID can act as a BAX-like effector of apoptosis. EMBO J. 41, e108690 (2022).

19. Denault, J.-B., Eckelman, B. P., Shin, H., Pop, C. & Salvesen, G. S. Caspase 3 attenuates XIAP (X-linked inhibitor of apoptosis protein)-mediated inhibition of caspase 9. Biochem. J. 405, 11–19 (2007).

20. Twiddy, D., Cohen, G. M., Macfarlane, M. & Cain, K. Caspase-7 is directly activated by the approximately 700-kDa apoptosome complex and is released as a stable XIAP-caspase-7 approximately 200-kDa complex. J. Biol. Chem. 281, 3876–3888 (2006).

21. Slee, E. A., Adrain, C. & Martin, S. J. Serial killers: ordering caspase activation events in apoptosis. Cell Death Differ. 6, 1067–1074 (1999).

22. McComb, S. et al. Efficient apoptosis requires feedback amplification of upstream apoptotic signals by effector caspase-3 or -7. Sci. Adv. 5, eaau9433 (2019).

23. Dumont, P. et al. The Amaryllidaceae isocarbostyril narciclasine induces apoptosis by activation of the death receptor and/or mitochondrial pathways in cancer cells but not in normal fibroblasts. Neoplasia 9, 766–776 (2007).

24. Miyoshi, N. et al. Alpha-tocopherol-mediated caspase-3 up-regulation enhances susceptibility to apoptotic stimuli. Biochem. Biophys. Res. Commun. 334, 466–473 (2005).

25. Zhu, J., May, S., Ulrich, C., Stockfleth, E. & Eberle, J. High ROS Production by Celecoxib and Enhanced Sensitivity for Death Ligand-Induced Apoptosis in Cutaneous SCC Cell Lines. Int. J. Mol. Sci. 22, 3622 (2021).

26. Schneider, C. A., Rasband, W. S. & Eliceiri, K. W. NIH Image to ImageJ: 25 years of image analysis. Nat. Methods 9, 671–675 (2012).

27. Marino, S., Hogue, I. B., Ray, C. J. & Kirschner, D. E. A methodology for performing global uncertainty and sensitivity analysis in systems biology. J. Theor. Biol. 254, 178–196 (2008).

28. Kennedy, J. Particle swarm optimization. in Encyclopedia of machine learning 760–766 (Springer, 2011).

29. Iadevaia, S., Lu, Y., Morales, F. C., Mills, G. B. & Ram, P. T. Identification of optimal drug combinations targeting cellular networks: Integrating phospho-proteomics and computational network analysis. Cancer Res. 10.1158/0008-5472.CAN-10-0460 (2010) doi:10.1158/0008-5472.CAN-10-0460.

30. Elmore, S. Apoptosis: A Review of Programmed Cell Death. Toxicol. Pathol. 35, 495–516 (2007).

31. Mustafa, M. et al. Apoptosis: A Comprehensive Overview of Signaling Pathways, Morphological Changes, and Physiological Significance and Therapeutic Implications. Cells 13, 1838 (2024).

32. Wolf, P. Tumor-Specific Induction of the Intrinsic Apoptotic Pathway-A New Therapeutic Option for Advanced Prostate Cancer? Front. Oncol. 9, 590 (2019).

33. Dhani, S., Zhao, Y. & Zhivotovsky, B. A long way to go: caspase inhibitors in clinical use. Cell Death Dis. 12, 949 (2021).

34. Tan, Y. S., Mhoumadi, Y. & Verma, C. S. Roles of computational modelling in understanding p53 structure, biology, and its therapeutic targeting. J. Mol. Cell Biol. 11, 306–316 (2019).

35. Wu, J., Haugk, K. & Plymate, S. R. Activation of pro-apoptotic p38-MAPK pathway in the prostate cancer cell line M12 expressing a truncated IGF-IR. Horm. Metab. Res. Horm. Stoffwechselforschung Horm. Metab. 35, 751–757 (2003).

36. Sun, X. et al. Systems modeling of anti-apoptotic pathways in prostate cancer: psychological stress triggers a synergism pattern switch in drug combination therapy. PLoS Comput. Biol. 9, e1003358 (2013).

37. Tian, X. et al. Targeting apoptotic pathways for cancer therapy. J. Clin. Invest. 134, e179570 (2024).

38. van Loo, G. et al. The role of mitochondrial factors in apoptosis: a Russian roulette with more than one bullet. Cell Death Differ. 9, 1031–1042 (2002).

39. Makinwa, Y., Luo, Y., Musich, P. R. & Zou, Y. Canonical and Noncanonical Functions of the BH3 Domain Protein Bid in Apoptosis, Oncogenesis, Cancer Therapeutics, and Aging. Cancers 16, 2199 (2024).

40. Zhu, X. et al. Autophagy activated by the c-Jun N-terminal kinase-mediated pathway protects human prostate cancer PC3 cells from celecoxib-induced apoptosis. Exp. Ther. Med. 13, 2348–2354 (2017).

