## Supplemental Figures for "A Mechanistic Model of Caspase-mediated Apoptosis Signaling Reveals Regulators of Drug Resistance in Prostate Cancer Cells"

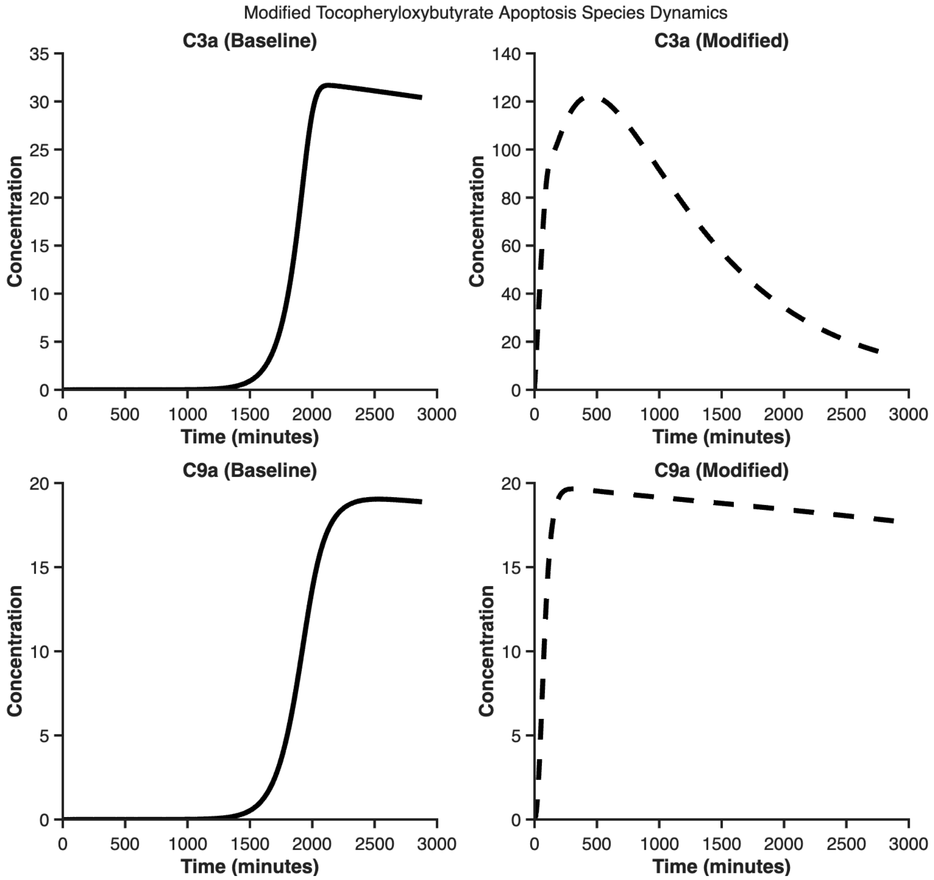


**Supplemental Figure 1: Modifications to produce a bell-shaped caspase-3 response.** The baseline model was modified to produce C3a and C9a dynamics qualitatively similar to data from the PC3 cell line treated with Tocopheryloxybutyrate treatment. In order to generate a bell-shaped caspase-3 response we (i) introduced $k_{Toco}$, which promotes more C3 activation by C8a, (ii) scaled the degradation of C3a by a sink parameter (β), and (iii) allowed $k_{Toco}$ to vary over time. Solid black lines, C3a and C9a dynamics produced by the baseline model derived from Harrington. Dashed black lines, dynamics simulated by the modified model.

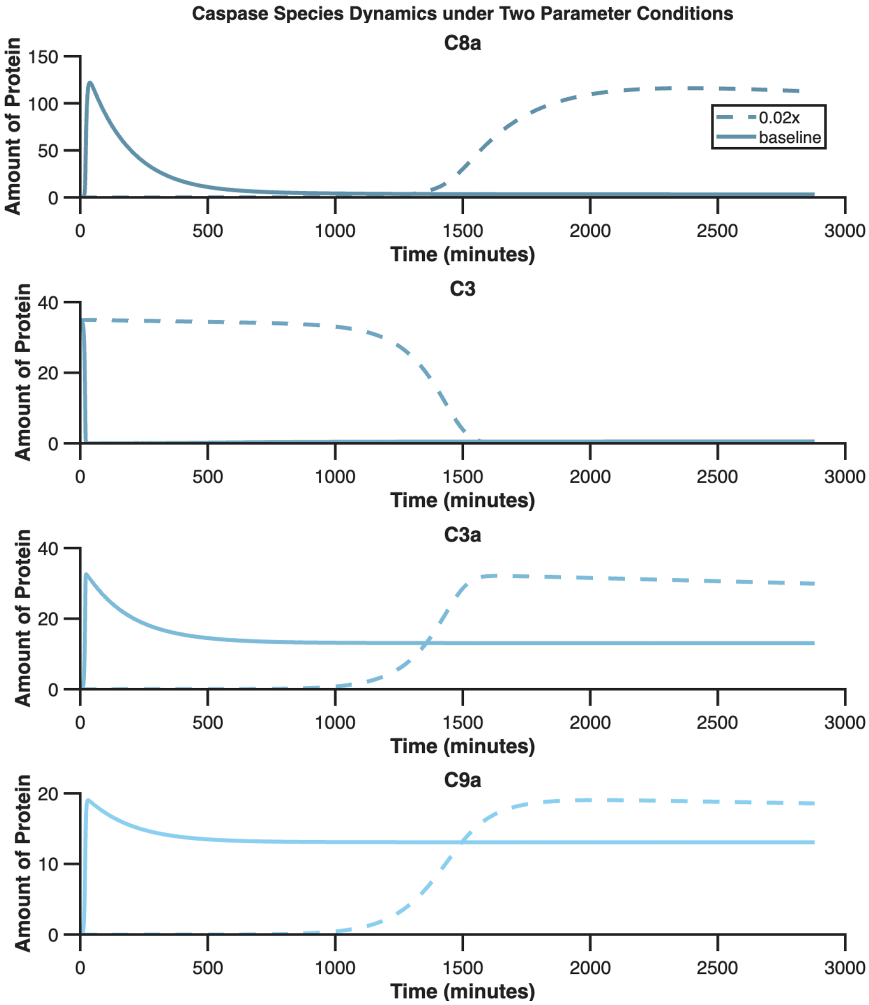


**Supplemental Figure 2: Comparison of dynamics for caspase activation.** Comparison between the baseline model from Harrington (solid line) and the modified model (dashed line), in which all reactions were scaled by 2% to better match the experimentally observed activation timing of caspases in the PC3 cell line.

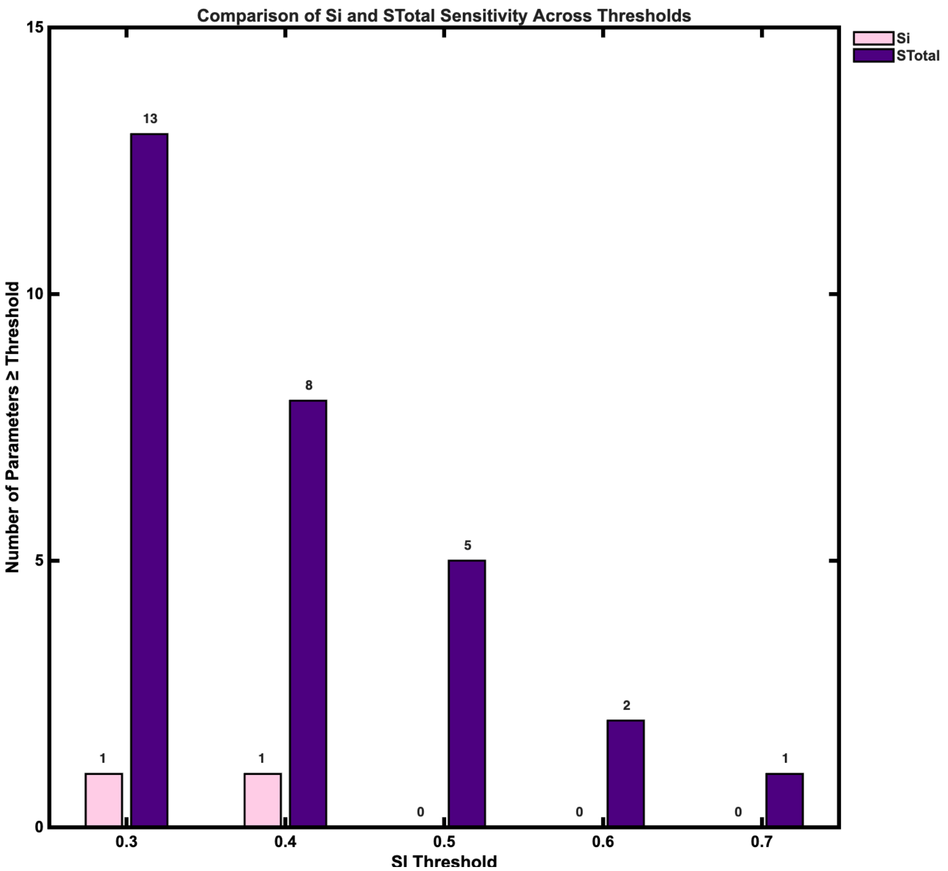


**Supplemental Figure 3: Number of parameters whose sensitivity indices reach a specified threshold**. Comparison of Si and STotal indices reveal that behavior is strongly influenced by nonlinear interactions between kinetic parameters and initial protein amounts, rather than isolated model inputs.

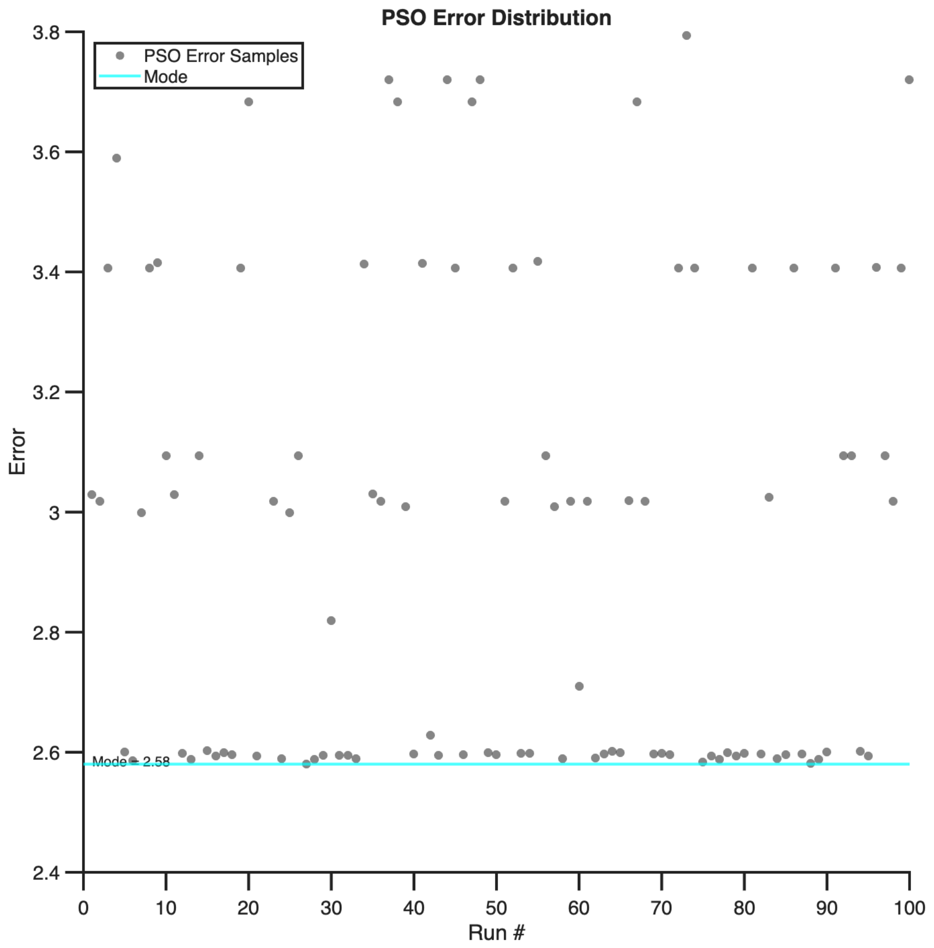


**Supplemental Figure 4: Error between model and data for simulations with estimate parameter values.** Distribution of SSE values corresponding to the 100 PSO runs shown in Figure 5. This illustrates the variability in model fit quality across parameter sets and confirms the presence of multiple comparably optimal solutions within the parameter space. Cyan line denotes the mode.

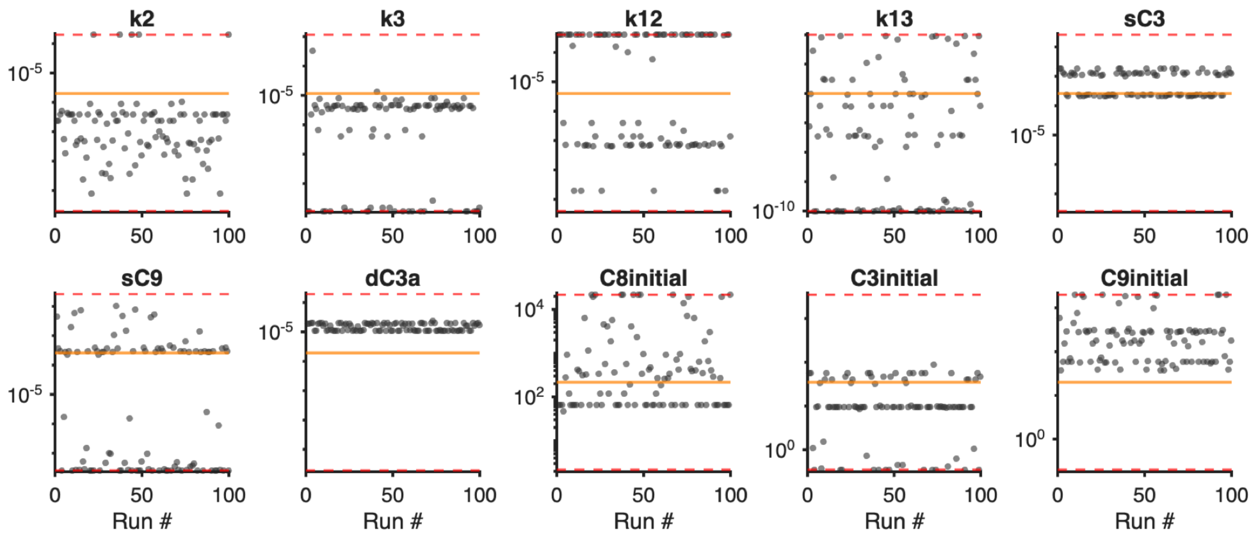


**Supplemental Figure 5: Estimated parameter values obtained using PSO.** Scatter plots show the distributions of best-fit values for the ten fitted model parameters across 100 optimization runs. Red dashed lines denote the minimum and maximum bounds imposed during optimization, yellow lines represent the nominal value, and each dot represents the estimated parameter from one PSO run.

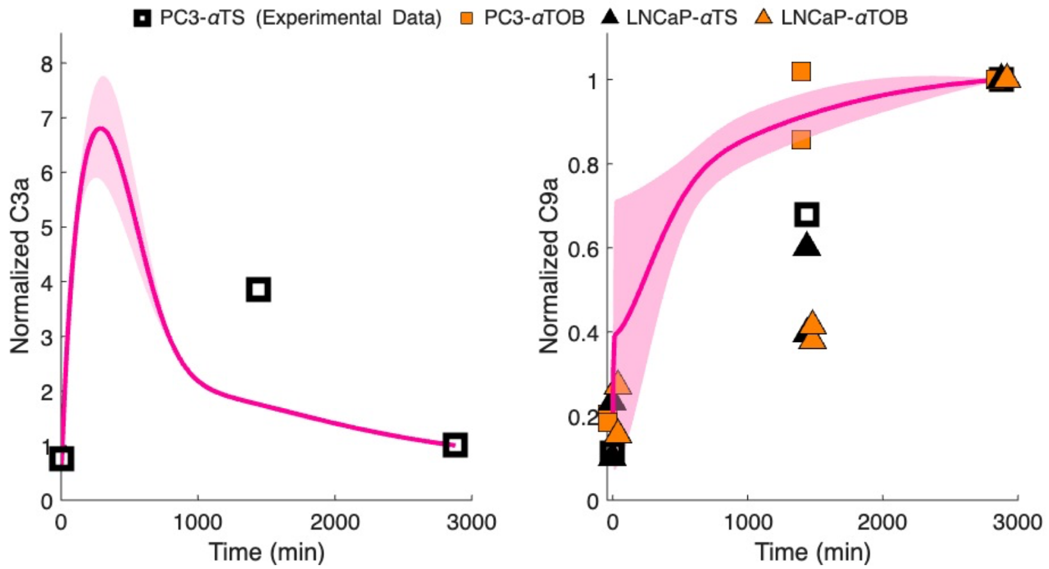


**Supplemental Figure 6: Comparison of model predictions to data not used in fitting.** Mean value for simulated dynamics for C3a and C9a using the fitted model are given in the solid line. Shaded regions represent the standard deviation across the best-fit parameter sets. Squares outlined in black are the experimental data used for model fitting, in the PC3 cell line with the αTS drug analog for Tocopheryloxybutyrate. We compare model simulations to additional data not used in fitting: the αTOB drug analog applied to the PC3 cell line (orange filled squares); αTS drug analog (gray filled triangles) and αTOB (orange filled triangles) applied to the LNCaP cell line.

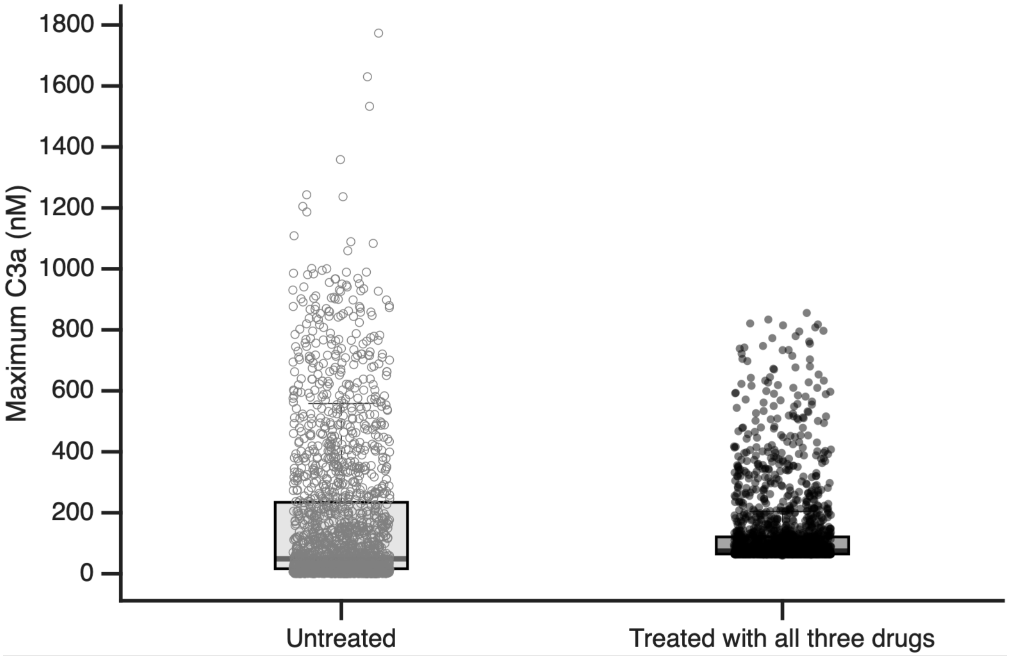


**Supplemental Figure 7: Effect of heterogeneity on maximum C3a.** Simulated results for the maximum C3a produced, with no treatment and with combination treatment (Tocopheryloxybutyrate, Narciclasine and Celecoxib). A heterogeneous population of 2,000 cells, each having a different initial amount of C8, C3, C9, BAR, and XIAP was simulated. Combination treatment increases the median maximum C3a from 48.9 nM (untreated) to 74.2 nM ($p$<0.01).

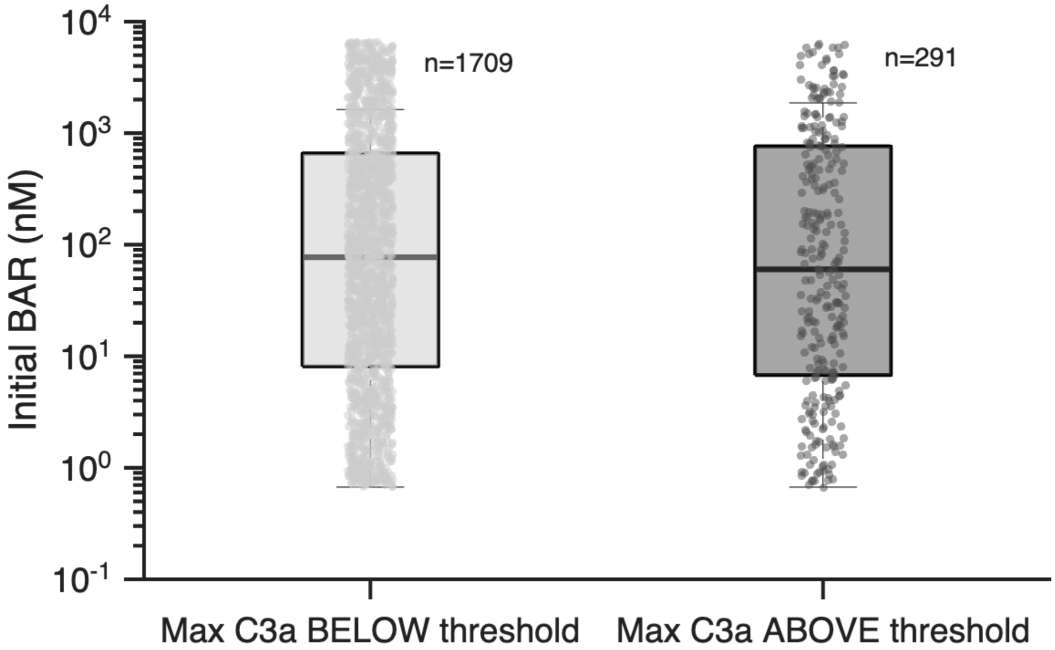


**Supplemental Figure 8: Effect of initial BAR concentration on maximum C3a produced.** Simulated results for the maximum C3a produced with combination treatment (Tocopheryloxybutyrate, Narciclasine and Celecoxib). A heterogeneous population of 2,000 cells, each having a different initial amount of C8, C3, C9, BAR, and XIAP was simulated. The initial amount of BAR is higher for cells in which the maximum C3a was below the threshold of 200 nM: 77.1 nM versus 60.3 nM, respectively ($p$<0.01).
